# Arousal-related mediation of perceptual belief updating across auditory domains

**DOI:** 10.1101/2025.09.10.675360

**Authors:** Roman Fleischmann, David Meijer, Burcu Bayram, Valentin Pellegrini, Ulrich Pomper, Michelle Spierings, Robert Baumgartner

## Abstract

Belief updating refers to the integration of prior beliefs with incoming evidence and guides decision-making under uncertainty. In response to surprising events, this process is thought to be modulated by the locus-coeruleus-noradrenaline (LC-NA) arousal system, observable via pupil dilations (PDs). Pertinent literature has mostly focused on conscious, high-level decision-making and estimation processes, while assuming that the same principles apply to low-level sensory perceptual decision-making and generalize across tasks, domains and modalities. To address some of these assumptions, we devised a novel perceptual discrimination paradigm, investigating behavior and PDs across auditory domains. Participants were presented with auditory sequences of randomized length at a rapid pace, changing intermittently between two latent states: acceleration vs. deceleration (temporal group, *N* = 25) and clockwise vs. counterclockwise movement (spatial group, *N =* 22). Under high uncertainty, participants continuously inferred latent states to report the final state per sequence. To extract per-stimulus estimates of PDs during sequences, we fitted a deconvolution-based general linear model to the continuous pupil traces with a free amplitude parameter reflecting ongoing PDs. A Bayesian observer model was fitted to participants’ responses and used to estimate information gain and surprisal for every stimulus. Participants’ performance and PDs showed strong sensitivity to the occurrence of change-points. Both computational variables significantly predicted PDs in both domains, with information gain outperforming surprisal in a model comparison. Further model comparison revealed significant preference for models excluding possible domain-specific effects over models including them, pointing towards a constant effect over domains. We conclude that behavior and associated PDs observed in our purely perceptual auditory task align with Bayesian principles of belief updating. The observed lack of domain specificity supports the assumed generalizability of belief updating.

## 1. Introduction

Belief updating describes a core mechanism of the brain. In theory, top-down expectations dynamically leverage learned regularities—beliefs—in order to suppress noise and enable predictions, while bottom-up sensory evidence enables adaptation to changes of circumstances and unexpected events, guiding cognitive decision-making in uncertain conditions [1], [2], [3]. For such a mechanism to drive sensory perception rather than higher-level cognitive decision-making, it needs to occur automatically, rather than consciously controlled. To drive perception on a universal level rather than under specific circumstances it further needs to generalize across domains and modalities, rather than occur in isolated tasks. The goal of this study is to investigate the influence of prior beliefs on decision-making under uncertainty on a generalized perceptual level.

Belief updating is mostly conceptualized within the Bayesian brain hypothesis [1] and the free energy principle [4], describing internal beliefs as *priors* which are integrated with incoming information— *likelihoods*—after weighing by the relative uncertainties [1], [2], [3], driving both perception and learning [4], [5], [6], [7], [8], [9]. The influential paradigm has early on been suggested to unify perception, learning, action and attention in a single model, to the point of offering a possible unified brain theory [1], [4], [5], [6], [8], [9], [10]. This proximity of perception and learning through a shared computational basis paired with the history of grand claims in the field has—between the lines—led to the somewhat liberal assumption that findings will translate between fields and Bayesian principles of belief updating will generalize across tasks, conditions, domains and modalities.

Belief updating in the specific context of perception based on (near-)Bayesian computational models has been applied to individual perceptual tasks across different modalities [2], [5], [11], [12], [13], [14], with early research predominantly stemming from visual experiments [15], [16], [17] and more recent studies also focusing on auditory paradigms [18], [19], [20], [21], [22]. Despite either explicitly targeting perception or placing findings in the broader context of perception, the used experimental task-designs tend to get conflated with more explicit estimation processes or learning-like elements requiring some degree of (meta-)cognition [20], [23], [24], [25]. Investigations of belief updating are per definition interested in predictions and current beliefs by participants. To assess them as directly as possible, experimental designs tend to explicitly ask for predictions based on past evidence [19], [20], [21], [26] or estimations in form of probabilities [27]. This inevitably induces consciously controlled cognitive processing into the tasks which arguably exceeds the automatic nature required for perceptual belief updating. Further common approaches artificially insert task-irrelevant oddballs to increase uncertainty which participants need to explicitly differentiate from similarly unexpected—but task-relevant— stimuli indicating a change of context, or just regular outliers within a current context [18], [19], [23], [28]. Such conscious categorization requires participants to reflect on the task’s probabilistic meta-structure: the probabilities associated with the occurrence of all task-relevant and irrelevant stimuli. This limits the interpretability for perceptual systems.

In a similar fashion, the issue of generalization across domains and modalities lacks the appropriate experimental evidence. Literature about belief updating in perceptual inference often implies it to be a higher cognitive process affecting all perceptual processes alike [3], [11], [13], [29]. While the pertinent literature has successfully provided experimental evidence spanning a variety of tasks grounded in a shared computational basis [3], [29], [30], each experiment represents isolated evidence for a specific task. Studies targeting the assumed generalization across domains or modalities are rare [31].

Investigating belief updating with a focus on its generalization opens up the question of a mechanistic link capable of spanning across perception and learning, domains and modalities. The locus coeruleus (LC) noradrenaline (NA) arousal system is often suggested as a possible modulator, offering a convenient explanation as arousal is known to affect learning, perception, action and attention alike [32]. Various theories have tied increased activity in the LC-NA arousal system to an increased perceptual sensitivity to novel stimuli [32], [33], [34], [35]. The LC-NA system is suggested to increase the gain of task-relevant stimuli [32], [36] and trigger a reset of neural networks to favor novel stimuli at the cost of ongoing processing [37], driving rapid adaptation to changing environments [20], [38]. In the context of Bayesian inference, a common interpretation is that prediction errors trigger an arousal response [19], [20], [39], which in turn suppresses prior beliefs and lends relative weight to sensory input [21], [38], [40], [41]. Accordingly, surprisal, a precision-weighted quantification of the prediction error, correlates positively with task-evoked pupil dilations [19], [38], [42]. Pupil dilations are in comparable literature most often seen as a proxy for LC-NA arousal [19], [20], [38]. It drives pupil dilation by releasing NA [32], [43], leading some authors to refer to the system as *pupil-linked arousal* system [20], [44].

The present preregistered [45] study investigates the role of pupil-linked LC-NA arousal in ongoing auditory perceptual belief updating under uncertainty. It relates pupil dilation to latent variables of perceptual belief updating, such as *surprisal*, estimated by fitting a Bayesian observer model. It further investigates participants’ performance as a complementary behavioral indicator of perceptual belief updating.

To this end, we devised a novel experimental paradigm asking two groups of participants to discriminate latent states, which are inferred from sequences of sounds in a change-point paradigm (Fig. 1). Latent states entail the relevant variables that are causal to the observations, requiring the participants to infer said latent states from the observations [46]. Per domain and group two diametrically opposite latent states change within a changepoint paradigm [47] over sequences of sounds: Acceleration vs. deceleration (temporal task, *N* = 24) and clockwise vs. counterclockwise movement (spatial task, *N* = 22), respectively. Participants were tasked with discriminating the last latent state of a sequence with random length and potentially multiple change points. To ensure automatic perceptual belief updating the experiment only asks participants for their percept at the end of every sequence in a two-alternative forced-choice design, refraining from asking explicitly for predictions. Crucially, the task is constructed such that participants are not required to keep track of the probabilistic meta-structure of the task, as each presented point of evidence (POE) carries all information to solve the task unambiguously. Contrary to other paradigms [19], [21], [23] the necessary high level of uncertainty is not introduced through the probabilistic meta-structure, but by varying evidence strength at the stimulus level, including very small changes (weak POEs) that were difficult to correctly perceive independently of context. To nevertheless stimulate integration over multiple sounds, a single POE is presented through more than one successive sound, necessitating involvement of working memory. To further ensure assessment of automatic belief updating, stimuli were presented at a rapid pace and in an uninterrupted fashion and all measures were quantified per sound within ongoing sequences.

**Figure 1.**
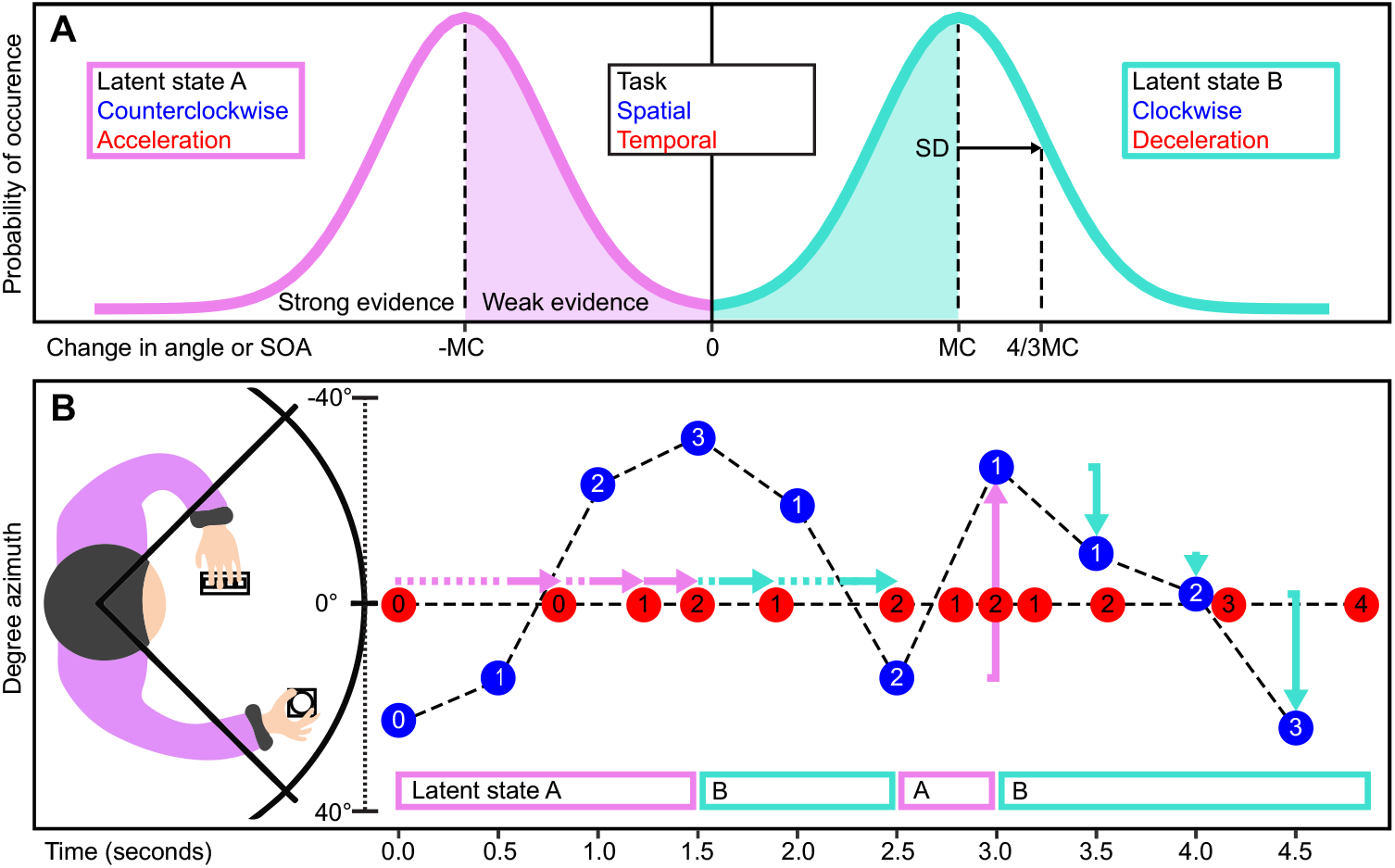
Task-schematic and sampling of points of evidence (POEs) *Note*. (A) Points of evidence (POEs) were sampled from two Gaussian distributions representing two distinct latent states: “acceleration” and “deceleration” in the temporal task, “clockwise” and “counterclockwise motion” in the spatial task. POEs are defined as changes in timing (horizontal arrows) or location (vertical arrows) of the current sound relative to the preceding sound. Changes in timing with constant location (dichotic listening) are POEs for temporal latent states, changes in location with constant timing are POEs for spatial latent states. The distribution and therefore the task’s difficulty was individualized via the mean change (MC). The shaded region between MC and -MC is considered to produce weaker POEs, vice versa the unshaded outside region is considered to produce stronger POEs. (B) Participants were presented with sequences of POEs of random length, interleaved with change points (CPs). Sounds were pink noise bursts defined by their location (angle in degrees azimuth) and stimulus onset asynchrony (SOA), shown as blue dots (±40°, SOA = 500 ms) and red dots (0°, mean SOA = 593 ms) representing two exemplary trials of the respective task-variation with three CPs each. Sounds are shown with their respective sound after change point (SAC) level, signifying the number of completed POEs within the current latent state, resetting at each latent state change (change point). Participants completed either the temporal or the spatial task variation. After each trial response confidence was assessed via 4-point Likert scale (“Very uncertain”, “Somewhat uncertain”, “Somewhat certain”, “Very certain”; left hand). During the task pupil dilation was measured via a desktop mounted eye tracker (not shown).

This leads to two main hypotheses: First, (H1) S*urprisal* is expected to positively correlate with ongoing pupil dilation on a single sound level and participants’ performance is expected to be biased by prior beliefs, impacting accuracy in relation to changes of latent state. Second, the experiment specifically tests across the spatial and temporal auditory domain, to distinguish a possible generalized modulation by LC-NA arousal from domain-specific effects. Hence, (H2) observed relationships between surprisal and pupil dilation are expected to be independent of perceptual domain, therefore comparable between tasks. In a further exploratory analysis we have tested how information gain (infogain), a latent variable closely related to surprisal [48], performs as a predictor of ongoing pupil dilation.

## 2. Results

### 2.1. Task performance and confidence increase with sound after changepoint (SAC) level

Average task accuracy binned per SAC level showed a significant main effect of SAC level (*F*_1.45, 63.88_ = 126.81, *p*_*GG*_ < .001, partial η^2^ = 0.742, 95% CI: [0.502, 0.960]) in a mixed-design repeated measures ANOVA, indicating that accuracy varied significantly as a result of the sequential presentation of POEs (Fig. 2A). In both tasks participants’ performance increased from below chance level to near-ceiling, with accumulation of consecutive coherent evidence for the current latent state after occurrence of a CP. A stark contrast is seen between SAC = 1 and later SAC levels. The largest increase in both tasks was observed from SAC = 1 to SAC = 2, also marking the crossing of chance performance at 0.5. Participants performed well below chance level for SAC = 1, translating to a tendency to answer according to the latent state from before the CP, rather than the presented sensory evidence. From SAC = 2 onwards, participants performed well above chance level. From SAC = 2 until SAC = 5 the increase in accuracy slows down in both tasks, although this pattern is more pronounced in the spatial task. Overall lower accuracies are found in the temporal task, a difference that proved significant as a main effect of domain (*F*_1, 44_ = 10.77, *p* = .002, partial η^2^ = 0.514, 95% CI [0.270, 0.898]). The interaction between SAC level and domain was not statistically significant (*F*_1.45, 63.88_ = 1.73, *p*_*GG*_ = .19).

**Figure 2.**
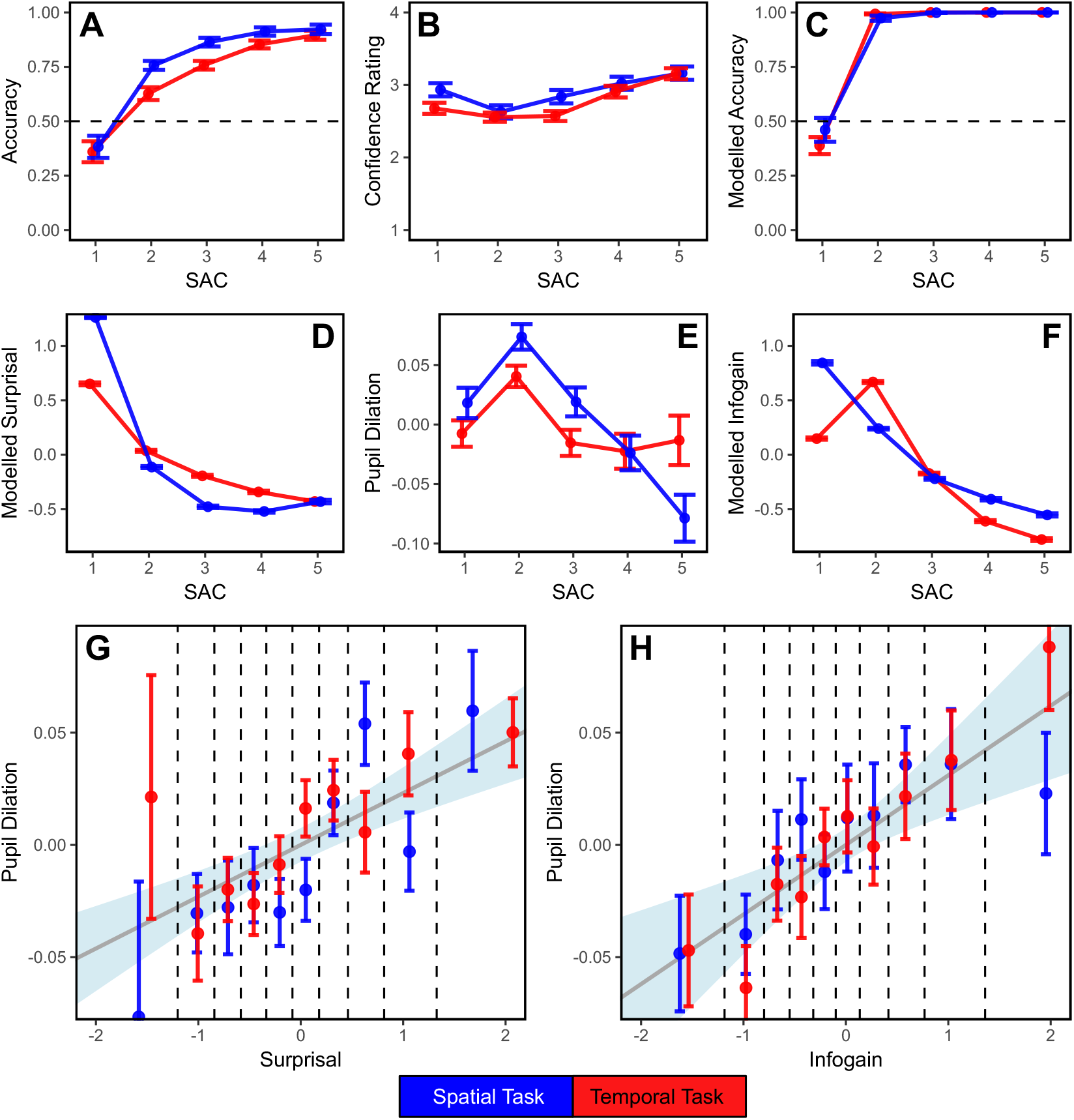
Measures of behavior and belief updating in relation to sequential presentation and pupil dilation. *Note*. (A) Group-average accuracy (percentage of correct trials) with chance level (dashed), (B) Group-average confidence ratings, (C) model predicted group-average accuracy (percentage of correct trials) with chance level (dashed), (D) model-estimated group-average surprisal, (E) group-average pupil dilation and (F) model-estimated group-average infogain. (A-F) All variables given as a function of sound after change point (SAC) level. Variables are given per trial (A-C) or per stimulus (D-F) respectively. (G) Pupil dilation as a function of surprisal and (H) infogain, binned in deciles and visualized with boundaries (vertical dashed). The slopes (grey line, ±95% CI in shade) reflect the relationships as predicted by winning mixed-regression models across both tasks. (D-H) Surprisal, pupil dilation and infogain were Box-Cox transformed and z-scored within participant. (A-H) Error bars denote group-level standard error of means across all panels.

Corresponding confidence ratings showed systematic variability over SAC level as well (*F*_2.54, 111.77_ = 46.54, *p*_*GG*_ < .001, partial η^2^ = 0.514, 95% CI [0.270, 0.898]), with a minimum at SAC = 2 (Fig. 2B). Participants indicated the least confidence in their choice at the same point within the sequence their average performance first crossed above chance level. Onwards from SAC = 2, with the increase in accuracy, confidence increased as well. Immediately after occurrence of a CP at SAC = 1, where below chance level performance indicates a tendency to the previous latent state, confidence ratings are low to moderate. The pattern of confidence shows lower ratings in the temporal task from SAC = 1 to SAC = 4, resulting in a prolonged, shallower negative peak compared to the sharp and defined negative peak with following linear increase in the spatial task.

From a Bayesian perspective, the observed patterns in accuracy and confidence are consistent with the influence of a prior belief continuously updated throughout the sequence. At SAC = 1, the first POE after a CP is presented and integrated with a previous (opposite and therefore detrimental) prior. This leads to consistently wrong answers under medium confidence. Only after the prior has been sufficiently updated through evidence under the new latent state—at SAC = 2—do answers become mostly correct, albeit with low accuracy and confidence. As consecutive POEs for the current latent state are accumulated without interruption—at SAC = 3 and beyond—the prior increasingly stabilizes perception, leading to improved accuracy and confidence.

### 2.2. Bayesian observer model and surprisal

The Bayesian observer model predicted participants’ answers in the spatial task with an average (predictive) accuracy of 84%; this corresponds to a mean *Cohen’s h* of 0.76 (STD = 0.17). In the temporal task the model performed at an average accuracy of 74%, which corresponds to a mean *Cohen’s h* of 0.51 (STD = 0.17). The model-based predictions of accuracy feature the same prominent minimum at SAC = 1, although predicting steadily high accuracies for all further SAC levels (Fig. 2C), thus outperforming participants’ accuracy. This is likely attributed to idealized and incomplete assumptions within the model. Participants are unlikely to integrate sequential evidence perfectly or acquire perfect knowledge of the generative process, as assumed by the model. Further, there might be additional sources of noise beyond the fitted sensory noise the model accounts for. This additional uncertainty likely leads to imperfect inference in participants. Nevertheless, the current—non-optimized—model proves useful in providing approximations of latent variables.

Model-based surprisal as a function of SAC level roughly resembles an inverse relationship to the observed average accuracy (Fig. 2D), mirroring both direction and magnitude of change. SAC = 1 again represents a stark contrast: as the POEs presented right after a CP represent a strong deviation to POEs presented until occurrence of a CP under the previous latent state, surprisal peaks at SAC = 1. Surprisal then decreases from SAC = 1 until SAC = 5, suggesting predictions to increasingly match the POEs. The largest decrease is observed from SAC = 1 to SAC = 2. After SAC = 2, decrease in surprisal is slowing down in both tasks, further underlining the stark outlier position of SAC = 1. Surprisal from the spatial task features overall more extreme values compared to surprisal from the temporal task; it is higher at SAC = 1, followed by a larger decrease and subsequent smaller surprisal at higher SAC levels.

### 2.3. Pupil dilation and pupil dilation rate are sensitive to change points and predicted by surprisal

Pupil dilations during the tasks’ sequences—extracted from a deconvolution-based model of the ongoing pupil signals on a per sound basis (excluding the first and last sounds in each trial)—show a systematic variability as a function of SAC (Fig. 2E), just as participants’ performance and surprisal. Pupil dilation therefore seems sensitive to the (detected) occurrence of CPs. In both domains, pupil dilation peaks at SAC = 2. This is the point of lowest reported confidence and the first point of performance above chance level. Subsequently, pupil dilation decreases past SAC = 2.

This observed pupil dilation pattern translates into a significant positive relationship of pupil dilation with surprisal across both tasks (Estimate = 0.023, SE = 0.004, *t*_41.76_ = 5.21, *p* < 0.001, n = 67,951 observations from 46 participants, R^2^m = .0005, R^2^c = .0008 in a mixed-effects linear regression model analysis). This shows that pupil dilations during the sequential presentation of POEs are predicted by model-derived estimates of surprisal across the temporal and spatial domains (Fig. 2G).

As indicated by the correlation of modelled pupil dilation with surprisal, the minimally processed (simple average across trials) pupil dilation rates (Fig. 3) also show a significant response to surprisal from 571 – 1249 ms, when comparing average pupil dilation rates for binned low-surprisal and high-surprisal conditions in a cluster-based permutation analysis (*p* < 0.001, summed cluster *t* = –2107.33). After a common baseline with two consecutive low-surprisal sounds, the averaged pupil size trace positively deflects in response to a condition-defining third high-surprisal stimulus, compared to occurrence of a third consecutive low-surprisal stimulus (0 ms, Fig. 3A). After deviation the pupil size traces resume in parallel. Since the total distance between the parallel pupil size traces after 0 ms is affected by the choice of baselining period, we performed the main analysis on instantaneous pupil dilation rates (Fig. 3B). Overall higher values in the “high surprisal” condition during the significant cluster mark a relatively stronger increase in the original pupil size traces, compared to the “low surprise” condition, as visible in Fig. 3A. Note that this analysis could only be performed on the data from the spatial task, due to its regular SOA, which allowed to superimpose a large number of trials with synchronized stimulus times.

**Figure 3.**
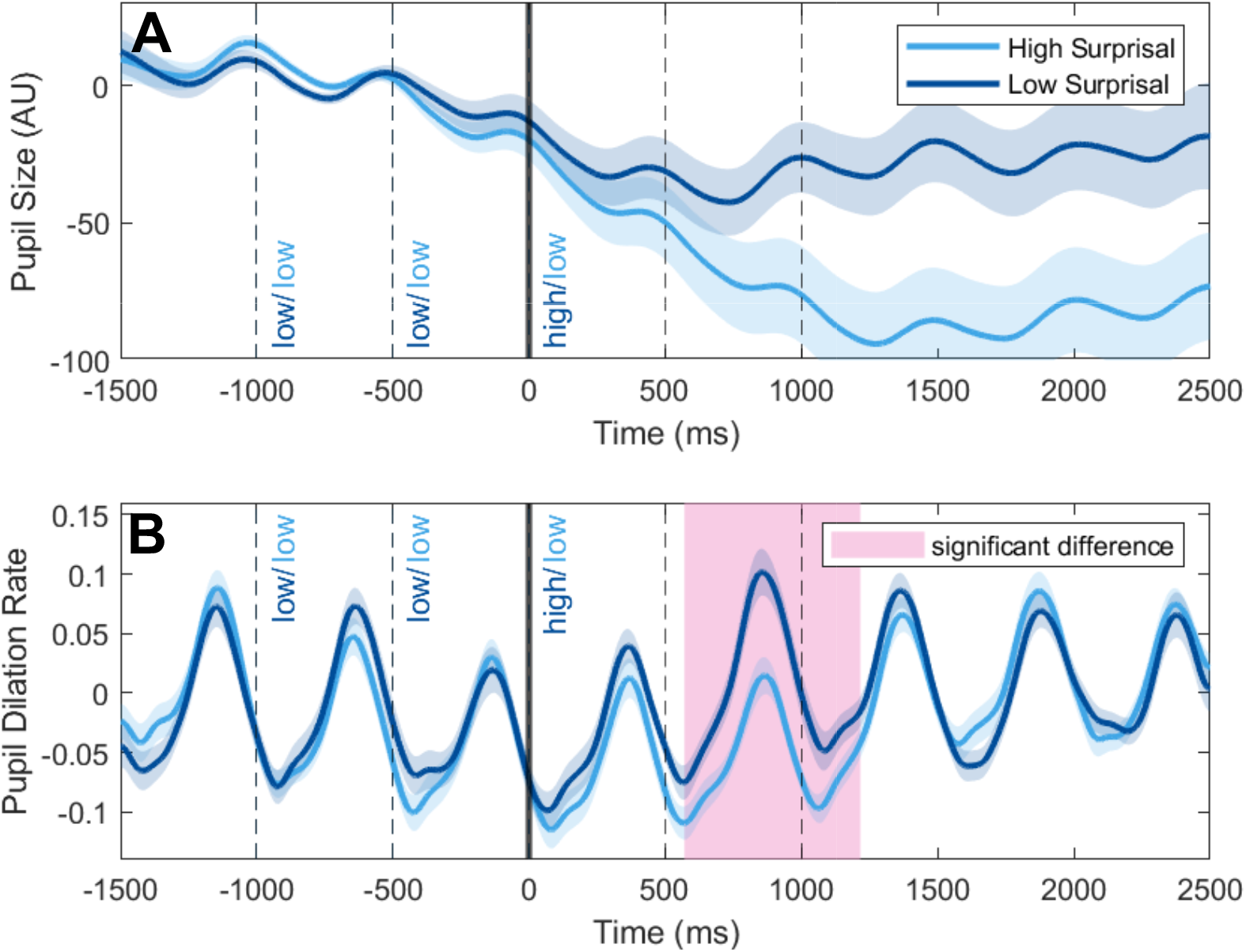
Effect of surprisal on pupil size and pupil dilation rate in the spatial task. *Note*. Average pupil traces binned into a “low” and a “high” surprisal condition. Two initial low-surprisal sounds (–1000 ms, –500 ms) are followed by a defining third stimulus (0 ms), either low or high in surprisal (median split per subject). (A) Pupil size traces deviate in conditions after occurrence of the defining stimulus. (B) Pupil dilation rates show the time period where the original pupil size traces deviate, marked in by overall higher values in the “high” condition. Both pupil size traces and dilation rate are shown with group-level standard error of mean (shaded area).

Fig. 3A also shows an overall decrease of pupil size after each low surprisal stimulus. The average pupil size trace of the low-surprisal condition strongly declines with three consecutive low surprisal sounds. The average pupil size trace of the high-surprisal condition initially declines in parallel with the low-surprisal condition for the initial two consecutive low surprisal sounds, then stops its decline after the defining third stimulus. Relative to the low-surprisal condition, this poses the positive deflection which is captured by the dilation rate-based analysis. The overall decline could be attributed to the chosen baseline period, which is entirely comprised of low-surprisal sounds even in the high-surprisal condition. Two consecutive low-surprisal sounds possibly lead (to an extend) to a constriction response or a return to baseline within an overall uncertain experimental paradigm, which cannot be reliably reversed by a single above-median surprisal sound.

### 2.4. Influence of domain on pupil dilation is negligible

A more extensive mixed-effects regression model including a fixed effect of domain and its interaction with surprisal on pupil dilation showed no significant improvement over the presented reduced model (*χ*^2^(2) = 0.63, *p* = 0.730) and model indices favored the reduced model (BF = 49587.4, ΔAIC = –4, ΔBIC = –21, ΔlogLik = 0). The fixed effect of task and the interaction term thus do not significantly contribute to explaining the variance in pupil dilation. This shows that the observed prediction of pupil dilation by surprisal is largely independent of the respective presented task and therefore domain.

### 2.5. Exploratory analysis suggests infogain as superior predictor of pupil dilation

A direct comparison between the so far established winning model for surprisal and a corresponding model for infogain (Estimate = 0.031, SE = 0.008, *t*_46.21_ = 3.80, *p* < 0.001, n = 67,951 observations from 46 participants, R^2^m = .0010, R^2^c = .0033) favored infogain across all information criteria (BF = 6.608128e+25, AIC = –119, ΔBIC = –119, ΔlogLik = 59). Infogain hence proved to predict pupil dilation during the trials’ sequences better than surprisal. This result is underscored by the observed pattern of infogain as a function of SAC level (Fig. 2F). While surprisal has a clear peak at SAC = 1 and rapidly decreases thereafter, infogain only decreases gradually in the spatial task and actually peaks at SAC = 2 in the temporal task. This is similar to the pattern of pupil dilation (Fig. 2E) and unlike the pattern seen in surprisal (Fig. 2D). The consistent advantage across all goodness-of-fit metrics alongside improved effect sizes (ΔR^2^m = .0005, ΔR^2^c = 0.032) indicates that stimulus-to-stimulus infogain explains more variance in pupil dilation than surprisal.

To validate the used mixed-effect regression model for infogain, a more extensive model including a fixed effect of domain and its interaction – analog to the analysis already done for surprisal – was performed, showing no significant improvement (χ^2^(2) = 1.88, *p* = 0.391). The model fit indices favor the reduced infogain model (BF = 26598.59, ΔAIC = –3, ΔBIC = –20, ΔlogLik = 1). The fixed effect of task and the interaction term thus do not significantly contribute to explaining the variance in pupil dilation. Infogain therefore demonstrates the same non-specificity for domain already observed for surprisal in an analog model comparison.

Surprisal and infogain represent closely related but distinct and usually very intercorrelated constructs [25], [27], [49]. While surprisal is indicative of the “unlikeliness” of an event within a current prior belief, infogain represents the change of beliefs as a result of that event, with less likely events often, but not necessarily, leading to larger changes. As a practical difference within the presented paradigm: Surprisal is only meaningful to the task up to a certain point. While POEs can get theoretically indefinitely surprising based on their probabilities of occurrence, the gain of information is limited to the degree that the prior can be updated, essentially up to a *certain* latent state change. From this point onwards stronger surprisal does not translate to more infogain. Vice versa, when priors are ambiguous, even small surprisal can cause relatively large effects on infogain.

## 3. Discussion

In the present study, two groups of participants were presented with two variations of a latent state discrimination task, one in the auditory spatial domain and one in the auditory temporal domain, while measuring their pupil dilation. We investigated the behavioral performance and pupillometric measurements for signs of arousal-modulated pupil-linked belief updating on a perceptual level spanning across domains. We report three main findings: First, (H1) participants showed ongoing belief updating on an automatic perceptual level as evidenced by systematic sequential variation of ongoing pupil dilation, pupil dilation rate and behavioral discrimination accuracy. Sensitivity of pupil dilation and pupil dilation rate to estimated surprisal indicates involvement of the LC-NA arousal system. Second, (H2) the relationship between modelled latent variables of belief updating and ongoing pupil dilation is not significantly affected by the tested perceptual domains, pointing towards a generalized mechanism. Third, in an exploratory analysis, infogain proved to be a superior correlate for pupil dilation in comparison to surprisal.

### 3.1. Ongoing belief updating occurs on automatic perceptual level

In the present study, we observed belief updating to occur on a rapid perceptual level in an automatic fashion. Both behavioral performance and pupil dilations indicated ongoing belief updating during a novel task, designed to require only simple perceptual judgements. The task was deliberately chosen to (a) avoid reliance on monitoring the probabilistic meta-structure of sequences, (b) not require explicit predictions, and (c) present sounds in a rapid, uninterrupted stream.

Compared to earlier studies, the current paradigm minimizes the need for extended contextual monitoring or explicit reasoning about probabilistic meta-structure. A comparable study [20] introduced uncertainty via weak perceptual evidence, but also required participants to provide explicit predictions. Other studies induced uncertainty via oddball stimuli [19], [23], forcing participants to judge whether unexpected events indicated a changed state or were irrelevant outliers [18], [19], [23], [28]. Such designs engage higher-level reasoning processes, whereas the present task reduced uncertainty directly by varying evidence strength, including very small changes (weak POEs) that were perceptually difficult to detect regardless of context.

The pacing of the task further reinforced this focus on perceptual processing. While slower or interrupted paradigms can facilitate pupil baseline measures [20] or more frequent behavioral responses [19], [21], they also risk introducing cognitive reflection on consecutive single stimuli. By contrast, the current paradigm employed rapid and continuous presentation (average SOAs: 593 ms for the temporal task and 500 ms for the spatial task), eliciting a streaming sensation rather than discrete evaluation of individual sounds. This ensured that observed biases toward prior beliefs were rooted in perceptual processing rather than conflated with explicit cognitive biases or strategies.

Behavioral performance showed a distinctive pattern well explained by accumulation of evidence in perceptual priors. At change points, discrimination accuracy dropped below chance level, demonstrating that perception was momentarily dominated by prior belief. This effect was amplified by the high level of uncertainty intrinsic to the task. As evidence accumulated under the new latent state, uncertainty decreased, reflected in gradually increasing accuracy. Under Bayesian observer theory, participants are assumed to estimate and account for those associated uncertainties, adapting to volatile environments and noisy observations [1], [26], [30], consistent with the present findings.

Modelled pupil dilations provided a complementary neurophysiological correlate to these observations. A positive relationship between surprisal and pupil dilation has previously been observed in Bayesian inference tasks [44], [50], likely reflecting LC-NA arousal responses to prediction errors [40]. The presented findings align with these observations and extend them to the auditory modality, supporting the notion of probabilistic integration of prior beliefs with sensory evidence [19], [20], [21].

The minimally processed pupil dilation rate showed sensitivity to high-surprisal stimuli as well, consistent with prior research [26], [42], [44], [50]. The pupil response to surprisal occurred between 571 – 1249 ms after stimulus onset, which matches the expected psychosensory delay of around 1second [51], although slightly on the faster side. Reported latencies in the literature vary: arousal-related effects have been documented as early as 400 ms [52], peaks in response to salient auditory stimuli appear at about 900 ms [53], and responses to unexpected stimuli are typically reported within 500 – 1500 ms [42], [54] [44], [55]. Together, these findings confirm our observation of a relatively quick pupil response to surprisal.

Interestingly, both forms of analysis suggest that effects of surprisal on pupil dilation or pupil dilation rate are not exclusively showing increases of pupil size, as both measures showed constricting responses as well. Low-surprisal stimuli often elicited constriction or a return to baseline, which was interrupted by relative dilation in response to high-surprisal stimuli. A possible explanation lies in pupil baseline fluctuations, which are commonly linked to uncertainty [20], [26]. Because the present experimental paradigm was designed to maintain high overall uncertainty, participants may have been in a state of elevated baseline pupil dilation. Such a state could impose a ceiling effect, limiting the magnitude of further task-evoked dilation. Indeed, evoked pupil dilations are known to vary inversely with baseline size [51], [56], [57], with higher baselines—often associated with uncertainty—producing smaller relative responses. Studies targeting exploration-exploitation trade-offs have similarly linked elevated baselines to reduced evoked dilations [58], [59], [60].

Within the current experimental conditions, two to three consecutive low-surprisal sounds could well mark the beginning of a descent into more certain, more excitable lower baseline territory. However, comparable studies report baseline recovery only after up to 6 seconds post stimulus onset [44]. Due to the rapid nature of the present task, reliable baseline measures could only be obtained before the start of a sequence and therefore ongoing baselines could not be analyzed.

### 3.2. Relationship between pupil size and latent variables of belief updating is not significantly affected by perceptual domain

The prediction of ongoing pupil dilation by surprisal did not differ significantly between perceptual domains, consistent with the preregistered hypothesis [45]. Infogain showed the same lack of domain specificity, reinforcing the finding. The hitherto described relationship between either surprisal or infogain and pupil dilation was better described as a general effect across both experiments than as separate effects within spatial and temporal domains.

This has implications for the question of the generalizability of probabilistic perceptual inference. The pertinent literature adopted a view carefully suggesting (near-)Bayesian principles to dictate perception [3], [29], [61] and learning [46] across modalities, domains and a huge variety of tasks. Influential theories discuss Bayesian perceptual inference at length with little mention of specific domains or modalities, suggesting all (perceptual) information to be represented probabilistically [1]. Some go further, proposing that the ongoing creation of expectations is the brain’s main purpose, tying the Bayesian brain hypothesis across perception, learning and attention to a possible unified brain theory [4], [8], [10]. The addition of the LC-NA arousal system as a possible modulator of belief updating further strengthens the idea of a system-wide mechanism transcending modalities or domains: Arousal is known to affect attention and most higher cognitive processing [62] as well as behavior [58], perception, and memory [63]. The present results suggest that this implied notion of generalizability seems plausible within the scope of spatial and temporal auditory perception, as pupil-linked arousal effects on belief updating extended across both domains. Yet, single-domain or single-modality tasks are inherently insufficient in order to draw conclusions about a possible generalization. Therefore, more direct comparisons over modalities, domains and tasks are necessary to investigate the notion of generalizability in (auditory) perception.

It is also important to point out that a postulated generalized mechanism within the auditory modality is not necessarily to be expected and the purposefully chosen domains and tasks have the potential to yield differing results. The auditory modality is seen to be the ‘appropriate’ one for the temporal domain, generally characterized to be highly sensitive and best equipped to detect temporal changes [64], [65], [66]. This stands in stark contrast to auditory capabilities in the spatial domain, which is best perceived in foveal vision [66], [67]. This pairing translates to a neural level, where spatial processing co-localizes with visual processing and temporal processing co-localizes with auditory processing. This specificity is further underlined as under high demand an ‘inappropriate’ modality for a task, e.g. the temporal modality for a localization task, will recruit from the other, in this example visual, attention network [66]. Within the auditory modality spatial and temporal tasks represent starkly differing demands for the perceptual and attention system. We interpret a possible bridging or unifying mechanism for belief updating—as observed via pupil dilation—to point towards a modulation of perception through arousal that transcends the tested domains.

### 3.3. Ongoing pupil dilation is better explained by infogain

Despite the preregistered focus on surprisal as a hypothesized correlate for pupil dilations, infogain was found to be a better predictor of pupil dilation, consistent with previous reports [55]. A computational study [49], re-analysing data from prominent studies [20], [50], poses the hypothesis that of the variety of processes and variables associated with pupil dilation, a large portion can be subsumed under—and explained by—infogain. Hence, the Kullback-Leibler divergence between prior and posterior could be the common denominator between results claiming a correlation between pupil dilation and mental effort, attention, surprisal, decision biases, uncertainty, exploration-exploitation trade-off and learning rate respectively.

The precise processes reflected in pupil dilations remain debated. While often interpreted as an index of arousal [57], [68], [69] (but see also: [39]), alternative accounts implicate neural gain [70], cognitive load [71], [72], or attentional states, alongside more peripheral influences such as luminance and visual focus [51]. Other neuromodulatory systems than LC-NA that are found to be connected to pupil dilations include acetylcholine [73], serotonin [74] and the hypothalamus orexin system [75]. In the context of belief updating, surprisal is most commonly suggested as a correlate for task-evoked pupil dilations [26], [42], but the pertinent literature also finds correlations to belief updating complexity [19], latent state transitions [76], [77], learning rate [78] and upcoming choice [56]. Studies further point towards uncertainty, although uncertainty is mostly suggested to be represented in slower baseline changes of pupil dilations [20], [26]. Any interpretation regarding pupil dilation therefore must be framed within this broader set of possible correlates.

It is crucial to further note that the presented task design cannot definitively disentangle surprisal and infogain beyond a model comparison, as the two measures are highly correlated and cannot be experimentally separated here. A recent series of EEG-studies [23], [24], [25] specifically targeted this issue. To overcome the problem of intercorrelation between suprisal and infogain, they color-coded stimuli to provide participants with cues about whether stimuli are indicative of a new latent state (change-point) or are just rare oddballs belonging to the existing latent state. Both types of stimuli elicit high surprisal, but only the former leads to an updating of beliefs (high infogain). They conclude that the two measures have meaningful differences within the process of belief updating. Further studies with similarly capable task designs are needed to investigate such differences on a pupillometric level.

## 4. Conclusions

The present study demonstrates that belief updating, previously mostly studied in the context of consciously controlled cognitive decision-making, also operates rapidly at a perceptual level. Behavioral and pupillometric results together support the view of ongoing sensory belief updating. Pupil dilation correlated with surprisal across both spatial and temporal auditory tasks, without domain-specific differences. This points towards a potential supra-domain modulation of perceptual inference by the LC-NA arousal system. Exploratory analyses revealed information gain to be a superior predictor of pupil dilation as compared to surprisal, suggesting that this variable may capture a more fundamental component of belief updating.

## Supporting information

Supplement: Bayesian Model Description

## Acknowledgements

This study was supported by an Austrian Science Fund (FWF) Young Independent Researcher Group (Grant-DOI: 10.55776/ZK66) to Michelle Spierings, Ulrich Pomper, and Robert Baumgartner. This project has received funding from the European Union’s Horizon Europe research and innovation programme under grant agreement No 101129903. Participant recruitment was supported by the Cognitive Science Hub of the University of Vienna

## 6. Methods

### 6.1. Participants

Two groups of participants were recruited for two variations of a task targeting either the temporal or spatial domain. After exclusions (see below), the groups consisted of 24 participants (temporal group, 11 female, *M* = 25.15 years, *SD* = 3.46 years) and 22 participants (spatial group, 12 female, *M* = 24.45 years, *SD* = 3.25 years) respectively. All participants reported to be healthy, possess normal hearing, have no neuronal disorders (like epilepsy or claustrophobia), have had no concussions, skull fractures, comas or operations on head or vascular system in the past, have taken no mood-altering medication (psychopharmaceuticals, hormones) and have not engaged in drug use on the day of testing and the day prior (including alcohol). Participants received monetary compensation (10€/h) and gave written consent after an initial introduction by the experimenter. After the experiment they were debriefed about the study’s purpose.

In the temporal group 10 out of 34 recruited participants were rejected or excluded prior or during analysis: five participants failed to reach at least 60% correct trials during training, one participant was excluded due to low performance (at chance level) in the main task, one participant ended the experiment due to personal reasons, one participant’s pupillometry data was corrupted due to a technical issue and two participants’ pupil data sets were considered unusable due to exclusion of >80% of relevant data (see: 6.7 preprocessing).

In the spatial group of 7 out of 29 recruited participants were rejected or excluded prior to analysis: five participants were excluded during pretests due to large estimations of minimal audible angles (see: 6.6), two participants dropped out between two sessions due to personal reasons.

### 6.2. Setup

Participants were seated at a distance of 75 cm from a computer monitor (48 x 27 cm, 1.280 x 1.024 pixels, refresh rate of 60 Hz) in a dimly lit, sound attenuated room. Their heads were supported by a chin rest, preventing motion and ensuring constant distance and eye-level. Participants’ left hand was placed on a keyboard to give confidence responses via four designated keys. Participants’ right hand was placed on either a rotating wheel to give answers on motion direction, or on arrow keys of a keyboard to give answers on tempo changes. Auditory stimuli were presented binaurally via in-ear headphones (ER-2, Etymotic Research, Inc., Grove Village, Illinois) at 48 kHz sampling rate. Each stimulus consisted of a pink noise burst (10 ms on/off raised cosine ramps, high-pass filtered from 250 Hz with 4th order Butterworth) presented at around 80 dB instantaneous peak SPL (65 dB short-time averaged; Bruel G Kjaer 2260 Investigator sound level meter). Sounds were 25 ms long in the temporal task and 50 ms long in the spatial task. Sounds were spatialized in the spatial task using individualized head-related transfer functions (HRTFs; see 6.6 for the measurement procedure). Throughout the experiment we measured pupil dilation using an eye tracker (EyeLink 1000 Plus; SR Research, Osgoode, Ontario, Canada) at a sampling rate of 1 kHz. We also monitored brain activity via a 128-electrode EEG (actiCAP with actiCH Brain Products GmbH, Gilching, Germany). The EEG data analysis is not part of the current study. The experiment was run on Windows 10 in MATLAB (The MathWorks Inc., 2021; version: 9.11.0 (R2021b), Natick, Massachusetts), using the Psychophysics Toolbox (Version 3.0.17)[79], [80].

### 6.3. Tasks

Two groups of participants were presented with a latent state discrimination task with 200 auditory sequences over the course of four blocks (50 per block). The sequences consisted of a series of distinct pink noise bursts representing continuously changing *points of evidence* (PE) for one of two distinct latent states, presented at a rapid pace of about two sounds per second. Depending on the group the latent state and task-variation was either *spatial*, referring to the two mutually exclusive states “clockwise motion” and “counterclockwise motion”, or *temporal*, referring to the two mutually exclusive states “acceleration” and “deceleration”. A single POE for the respective latent state is defined as a change in a sounds’ location or timing compared to the preceding sounds’ location or timing. Changes in timing with constant location (dichotic listening) are POEs for temporal latent states, changes in location with constant timing are POEs for spatial latent states. Note that in this sense “acceleration” and “deceleration” refers solely to the pace of presentation throughout a sequence in the sense of rhythmicity, not to the speed of spatial motions. Latent states changed sporadically, with sequences potentially containing multiple change points (CPs). Participants were tasked with discriminating the final latent state in a two-alternative forced-choice design, which required them to infer it from the presented POEs. To ensure constant attention, sequence lengths were pseudorandomized, making every sound potentially task-relevant. During trials participants were instructed to focus on a central fixation dot to minimize eye movement. After providing the latent state estimate, each trial ended with a response on confidence on a 4-point Likert scale (“Very uncertain”, “Somewhat uncertain”, “Somewhat certain”, “Very certain”, left hand in Fig. 1B).

### 6.4. Generation of sequences

POEs were sampled on a stimulus-to-stimulus basis in a two-step process: First a latent state is set by randomly choosing one for the first POE (p=0.5) and then changing it for each new POE with a hazard rate of *H* = 0.2. Then the change in timing or location is sampled from the respective normal distribution, which was individually centered around a mean change (MC, *SD =* MC / 3) per participant (see Fig. 1A) to personalize the task’s difficulty. Each completed POE had the same chance of being the final one (pseudo-randomly sampled for a geometric distribution, p = 0.1), giving sequences an average length of 10 POEs.

Here absolute changes in timing or location smaller than a person’s MC (the region between +MC and –MC) are considered weaker POEs for the current latent state (under the premise of no CP), which are making the inference and task harder. Vice versa absolute changes larger than |MC| (the regions outside +MC and –MC) are considered stronger POEs, making inference and task easier. The individual MC was chosen such that it represents a rather demanding baseline level of evidence for the respective participant aiming for 80% correct trials.

Note that the Gaussian distribution of the normal distribution was truncated: POEs that were sampled from the “other” side of the distribution (0.1%) which is not congruent with the current latent state were disregarded and resampled. The experiment therefore does not include oddballs, where a POE from a differing latent state is presented without it indicating a latent state change. The task is therefore theoretically always possible to solve by observing only the final evidence (disregarding internal sensory noise), without having to learn the specific stochastic properties of the sequences.

### 6.5 Specifications of the temporal task

The temporal task referred to the two distinct latent states *acceleration* and *deceleration*. To this end we manipulated only temporal properties, the stimulus onset asynchronies (SOAs), while keeping the spatial properties static (diotic listening). Evidence for the current latent state was defined as *change in SOA* relative to the preceding SOA and sampled for each stimulus from the side of the bimodal distribution that represented the current state. For example, if the current latent state was “acceleration”, POEs would be sampled from the left side (Fig. 1A) which translated to a negative change in SOA, reducing the SOAs for each consecutive sound, and vice versa. A CP translates to sampling from the other side of the bimodal distribution, switching between positive and negative changes.

Participants were tasked with reporting the last latent state via arrow keys on a keyboard (up = speed up, down = slow down) after the end of a sequence. Participants completed in total six blocks of 50 trials. The last four blocks were the main experimental blocks and only during those, participants’ pupil dilation and brain activity (EEG) was monitored. Participants were offered breaks after every block to take as needed. After 25 trials within each block, participants were given a short break of 20 s. The first two blocks were used as training for participants to learn the task and its stochastic properties, and to individualize task difficulty.

Difficulty was set via one central parameter defining the sampling distribution: the temporal mean change (MC_t_). The bimodal distribution was defined as a normal distribution capped and horizontally mirrored at 0, with a mean of M = MC_t_ and a standard deviation of SD = MC_t_/3. In the temporal task, change was defined as a fraction of the previous SOA, assuming constant Weber fractions.

During the first two blocks the task difficulty was individually dialed in, aiming at an accuracy of ∼80%. Initially MC_t_ was set to 32.5%. After every ten trials in the first two blocks, difficulty was adjusted based on performance: MC_t_ was decreased by 1.25 percentage points if more than eight trials were answered correctly and increased by the same amount if less than eight were correct. For exactly eight correct answers, MC_t_ remained unchanged. MC_t_ was not increased beyond 35%, giving MC_t_ a range of 20% – 35% over 10 reevaluations in 100 familiarization trials. Participants not achieving at least 60% correct trials during the first two blocks were excluded from the experiment.

To achieve the desired change in tempo the previous SOA would be multiplied or divided with the sampled change + 1 depending on the current latent state. Considering for instance a sampled change of 30%, an exemplary previous SOA of 500 ms would be multiplied by 1.3, resulting in a change of SOA of 150 ms and a new SOA of 650 ms, therefore deceleration. Vice versa division by 1.3 would result in a change of SOA of –115,4 ms (new SOA of 384,6 ms) and therefore acceleration. The smallest sampled change could only be 0% in either latent state resulting in the new SOA being equal to the previous SOA.

The first SOA of any sequence was sampled at random and sequence generation was controlled to not contain SOAs smaller than 200 ms or larger than 1333 ms. The initial latent state was randomly sampled with equal chance and changed with the occurrence of a CP at *H* = 0.2, resulting in a range of 0 – 13 CPs per trial, depending on trial length. Any stimulus past the third stimulus had a 10% probability of being the last one, resulting in sequence lengths from 3 to 46 sounds.

In the four main experimental blocks sequences were brute-forced (i.e., generated at random until they met imposed criteria by chance) for all trials of a block at once to ensure moderately fair distributions of total sequence lengths, SAC level after last CP, last latent states, last level of evidence (SOA change), last SOA, as well as average hazard rates between *H* = 0.19 and *H* = 0.21. During familiarization participants received color-coded feedback (green = correct, red = incorrect) for every trial and a total number of correct trials after each block. During the four main blocks they only received the summary after each block.

### 6.6. Specifications of the spatial task

The spatial task required to dissociate between the two distinct latent states of *clockwise* and *counterclockwise motion*. To this end we manipulated only spatial properties, the azimuth angle of the sounds, while keeping the temporal properties constant; i.e., the SOA was kept constant at 500 ms. Evidence for the current latent state was defined as *change in angle* between two consecutive sounds and sampled for each stimulus from the respective Gaussian distribution. E.g., if the current latent state was “counterclockwise”, POEs would be sampled from the distribution on the left side (Fig. 1A), translating to a negative change in degrees azimuth and resulting in a position further to the left relative to the previous stimulus, and vice versa. A CP again resulted in sampling from the other—essentially inverted—distribution, switching between positive and negative changes. Participants were tasked with reporting the last latent state via a rotating wheel after the end of a sequence. During piloting button presses were used instead, leading to a bias in participants answers towards the hemisphere of stimulus presentation. This seemed to be caused by the ambiguity of “left” and “right” buttons, which could be interpreted as both the direction of the presented motion or the location of a stimulus. Substituting “left” and “right” with “clockwise” and “counterclockwise” to be indicated with the turn of a wheel proved to be an unambiguous alternative, solving the issue of the observed bias. Participants completed a total of five blocks with 50 trials each of the full spatial task of which only the last four were performed while measuring pupil dilation and brain activity (EEG). The first block was considered training and was not analyzed. Additionally, participants performed two blocks of a reduced spatial task, the familiarization task (50 trials each), and between three and five blocks of a separate variant of the familiarization task designed to estimate the participants’ minimal audible angle (MAA), later referred to as the MAA task (100 trials each). The entire experiment was split in two sessions on two separate days with a maximum of 72 hours in between. The first session started with one block of familiarization, followed by all MAA blocks and ended with one block of the full spatial task for training. The second session started with a block of familiarization and was followed by the four main blocks of the full spatial task that formed our data set for analysis. Participants were offered breaks after every block to take as needed. Between block 2 and 3 of the main four testing blocks we scheduled a mandatory break of at least 10 minutes. Within each block we scheduled a 30 s break every 25 trials.

In the familiarization task a trial consisted of one motion presented as a sequence of two sounds: a “standard” sound and a “probe” sound. The standard sound was randomly presented as starting-or endpoint of the motion and always at 0° azimuth; therefore, the probe was always presented at a peripheral position in space. The change of angle between standard and probe per SOA is referred to as one motion. Note that the SOA is a constant in this task, unlike in the temporal task where it is continuously changing. The task started easy with a motion of ±20°/SOA, successively decreasing its absolute value by 1°/SOA for ten consecutive trials. This was followed by 40 trials, randomized between ±20°/SOA and ±1°/SOA. The latent state of motion direction was pseudo-randomized. The participants received color-coded feedback (correct = green, incorrect = red) after each trial.

The MAA task followed the same concept, presenting one motion comprised of standard and probe. The standard was set at fixed locations that changed through blocks: 0° in block 1, ±20° in block 2, and ±40° in block 3. In blocks 2 and 3 the standard was always presented on the same lateral side for 25 consecutive trials. The upcoming side was announced during the breaks. The participants again received feedback for every trial as in the familiarization task. To estimate an MAA, the absolute change of angle between standard and probe was adjusted adaptively. The motions (and therefore difficulties) of the trials were adaptively set by the Psi-marginal algorithm [81], [82], [83]; using code from https://github.com/lacerbi/psybayes). We obtained estimates of the minimal change of angle between two sounds that participants needed to correctly infer the direction. We refer to these estimates as MAAs. MAAs are obtained per participant for three eccentricities in the respective blocks: at 0°, ±20° and ±40° azimuth. MAAs in both hemispheres are assumed equal, so their results were averaged. If the participants exceeded 3.3° at 0° azimuth (MAA_0°_), or 10° at either ±20° or ±40° azimuth, they were offered to repeat the block, once for each eccentricity. If the MAA still exceeded the threshold after one repetition, the participant was excluded from the experiment. This was necessary, as MAAs set the basis for the individualization of task difficulty, with larger MAAs translating to an easier task (see: below). For MAAs larger than these thresholds it was impossible to generate trials with longer sequences without exceeding the bounds of space set to ±40°. Outside those bounds of space MAAs can increase rapidly (and non-linearly) [84].

The main spatial task presented participants with sequences of consecutive motions representing POEs for the current latent state. Like in the temporal task POEs for latent states, in this case angular location changes in a clockwise or counterclockwise direction, were sampled from the respective Gaussian distribution defined in the same way via a spatial mean change (MC_s_): a normal distribution capped at 0, with a mean of M = MC_s_ and a standard deviation of SD = MC_s_ /3 (see Fig. 1A). The unit of change in the spatial task was in degrees azimuth, defining each stimulus position relative to the position of the previous stimulus. The individualization of task difficulty was achieved by defining the MC_s_ through the estimated MAA at the midline position as MC_s_ = 3 x MAA_0°_. The first stimulus location was sampled at random within the bounds of space (±40°). The initial latent state was randomly sampled with an equal chance and changed with the occurrence of a CP at *H* = 0.2, resulting in a range of 0 – 13 CPs per trial, depending on length. Any stimulus past the second stimulus had a 10% probability of being the last one, resulting in sequence lengths from 2 to 45 sounds. Participants received no feedback on single trials, only the number of correct responses over the whole block. Sequences were brute-forced (i.e., generated at random until they meet imposed criteria by chance) for all trials of a block at once to ensure quasi-fair distributions of total sequence lengths, sequence lengths after last CP, last latent state per lateral space quintile (within ±40°), last level of evidence (velocity) per latent state and per lateral space quintile, as well as hazard rates between 0.1 and 0.3.

For stimulus spatialization, we recorded each participant’s HRTF. Participants were positioned at the center of a spherical loudspeaker array (E301, KEF, radius 1.2 m) within a semi-anechoic chamber (T60 = 50 ms). Small microphones (KE4-211-2, Sennheiser) were inserted into their ear canals to capture audio during 91 exponential sweeps (one per loudspeaker) from 20 Hz to 20 kHz over 6 seconds, multiplexed across directions. The recorded HRTFs were then postprocessed to remove any acoustic effects from the chamber and its setup by equalizing them with the transfer functions of the sphere’s center point (previously measured without a participant present) and windowing the impulse responses to 5 ms. Multiplexing and windowing techniques followed a previously outlined procedure [85]. To achieve a 1° azimuthal resolution, we spatially upsampled the HRTFs via vector base amplitude panning [86] while using a spherical head model to correct for the direction-dependent time-of-arrival [87], as explained further elsewhere [88]. This customized set of HRTFs was applied for individualized stimulus spatialization. However, for 6 of the 22 participants generic HRTFs from a KEMAR mannequin [89] had to be substituted because technical equipment issues prevented accurate HRTF recordings.

### 6.7. Preprocessing of pupil size traces

Pupil size data represents the pupil diameter and was recorded in arbitrary units (AU) based on the image pixels belonging to the pupil area, as provided by the eye tracker. Pupil size data was first cleaned of blinks, gaps and recording artifacts. Blinks were detected using the *detectBlink()* function from the PUPILs toolbox [90], defining velocities faster than three times the median root mean square value of all successive 500 ms windows over the entire trace as blinks and marking them as missing data. Additionally, samples whose recorded values were lower than 3200 AU were marked missing data. Missing data were linearly interpolated up to 500 ms to reconstruct the pupil trace, longer periods were excluded from analysis. In this process two entire data sets from the temporal task were excluded from further analysis as 80% of sounds were excluded due to excluded corresponding pupil traces. All remaining data sets remained well below a maximum of 70% excluded sounds (spatial task: M = 36%, SD = 13%, temporal task: M = 32%, SD = 11%). All processed and interpolated pupil size traces were visually inspected and compared to raw pupil data to double-check the preprocessing.

For the instantaneous pupil dilation rates a zero-phase low-pass finite impulse response (FIR) filter was applied to the pupil size traces to attenuate frequencies above 4 Hz. The filter was designed as a Hamming window (filter order = 100, 1 kHz sampling rate). To eliminate phase distortion, the filter was applied in both forward and reverse directions using MATLABs *filtfilt()* function. We then approximated the first derivative as the difference between adjacent samples to obtain the instantaneous pupil dilation rate. No filtering was applied to pupil size traces entered into the deconvolution-procedure (see 6.8).

### 6.8. Deconvolution-based pupil model

Due to the rapid time-scale of the experimental design and the sluggish nature of the pupillary response [91] the evoked pupillary responses are convoluted and not directly accessible or interpretable. To extract a pupil dilation per sound event a deconvolution-based general linear model was fitted to the continuous pupil dilation traces, additionally informed by stimulus onset timestamps, following Denison et al. [91]. This is essentially a regression analysis, where the columns of the design matrix consist of time-delayed templates of pupil response functions [53] according to each sound’s latency, and the fitted coefficients represent free amplitude parameters. The amplitude parameters were used as estimates of the sound-evoked component on a single sound level, throughout this manuscript referred to as *pupil dilation* and given per sound event. Note that pupil dilation is differentiated from the previously described minimally processed *pupil size* and *pupil dilation rate*, which are both model-free continuous signals. We excluded the first and last 1500 ms of each sequence from analysis, as the pupil model performs worse at the edges. All pupil dilations for sounds falling into excluded periods of the preprocessed or predicted pupil trace are excluded from analysis, which includes the first two sounds of every sequence. Pupil dilations corresponding to sound-onset times of an exemplary trial alongside the raw, preprocessed and predicted pupil traces can be seen in Fig. 4. Over each participant’s pupil traces and corresponding model-predicted pupil traces (without excluded periods) we calculated the coefficient of determination R^2^ to quantify the explained variance of the deconvolution-based general linear model. On average it explained 92% of the pupil trace’s variance (mean R^2^ = 0.92, STD = 0.02).

**Figure 4.**
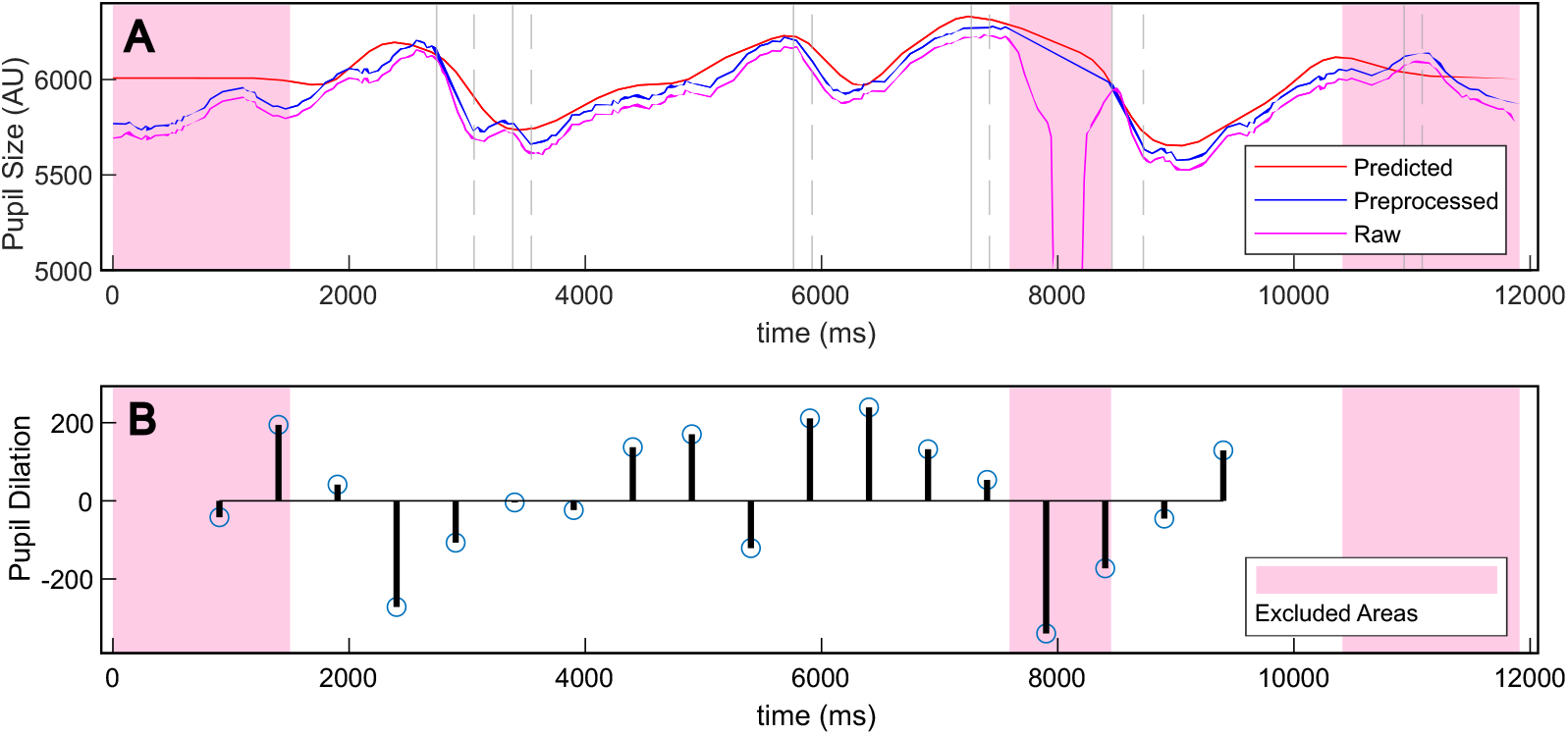
Pupil size traces, predicted pupil size and modelled pupil dilation for corresponding sounds. *Note*. (A) Unprocessed (raw), preprocessed and predicted pupil traces for an exemplary trial of the spatial task. Vertical lines mark start (solid line) and end (dotted line) of interpolation periods. Pink shaded areas mark periods and areas that are excluded, either a priori (first and last 1500 ms, values under 3200 AU) or as a result of blinks and artifacts that cannot be interpolated due to either a length of over 500 ms or a position towards the edges. (B) Modelled pupil dilation quantified by the amplitude parameter per sound of the same exemplary trial, with exclusion periods.

### 6.9. Bayesian observer model

A Bayesian observer model was fitted to the behavioral responses to compute momentary estimations of computational latent variables such as surprisal for every sound of every trial.

The observer model used in the current task assumes full knowledge of the sequence generation parameters, including the hazard rate (described in *supplements*) and estimate the other parameters that define the task-relevant uncertainty (prior on latent state *p* = 0.5, hazard rate *H* = 0.2, sampling distribution with *M* = MC, *SD* = MC/3, individually fitted sensory noise). We assume human observers to roughly equivalently learn the generative process during training blocks and use this knowledge of task-relevant uncertainties for perceptual inference. Participants were superficially informed by the experimenter about the concepts of latent states, CPs and varying evidence for latent states, but without mention of any specific values. To estimate sensory noise, the observer model adds random noise to each PE, blurring the previously unambiguous separation in two latent states and adding uncertainty. When receiving a noise-corrupted internal signal for the latest change of angle or SOA, the observer model then computes the relative likelihoods for the latent states that could have produced this internal signal. It then multiplies the likelihood ratio with the prior probability ratio for the two latent states to obtain a posterior probability ratio for the current latent state. This posterior ratio becomes the new prior ratio after adjustment for the possibility of a change point (and so further, until the end of the sequence). To account for sensory noise (which is inaccessible to the experimenter), the Monte-Carlo method was used to simulate 1000 trials with random-noise disturbed internal signals of SOA or angle change and compute estimations of latent variables as medians across all simulations.

For every stimulus in a sequence, surprisal can be computed based on likelihood ratio and prior ratio. We defined surprisal as the Shannon information, the negative log probability of a POE given the prior. Surprisal *S*_*t*_ at time point *t* is therefore computed based on the current priors *P*_*s,t*_ and likelihoods *L*_*s,t*_ per latent state s ∈ {*A, B*} as follows:

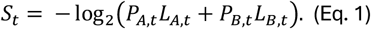

Across Monte Carlo simulations we calculated the median surprisal to obtain a single value per stimulus. For subsequent exploratory analyses we further quantified *information gain* (infogain). It is defined as the Jensen Shannon distance from prior to posterior and quantifies the gain of information as the result of the integration of the current stimulus. Infogain is thus defined as:

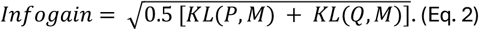

with *P* = {*Prior*_*A,t*_, *Prior*_*B,t*_} the prior probability distribution over the two latent states, *Q* = {*Post*_*A,t*_, *Post*_*B,t*_} the posterior distribution, *M* = {0.5(*Prior*_*A,t*_ + *Post*_*A,t*_), 0.5(*Prior*_*B,t*_ + *Post*_*B,t*_)} and the Kullback-Leibler Divergence denoted as 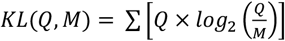. Infogain is closely related to what other literature often calls “belief updating” [22], [23], [24], [25] and sometimes “Bayesian surprise” [27], [92]. We chose the term infogain as a more descriptive variant and to avoid confusion of the estimated latent variable with the entire process of belief updating as described in the introduction. Due to inconsistencies of nomenclature it is advised to trace latent variables back to their computational definition when comparing surprisal and infogain. We included infogain due to the variety of literature targeting and comparing surprisal and infogain in the general context of perceptual inference [22], [23], [24], [25], [27] and tying infogain to pupil dilation [49], [55], [93].

### 6.10. Statistical analysis

#### 6.10.1. Variables and transformations

Per sound in every sequence, one estimate of pupil dilation was obtained from the deconvolution model (as described in section 6.8) and one estimate of surprisal was obtained from the Bayesian model (as described in section 6.9). To stabilize the variance, variables were Box-Cox-transformed within each participant and then standardized by z-scoring. Furthermore, each sound (and therefore every estimate of surprisal and pupil gain) has an assigned SAC level (Fig. 1). To only analyze POEs occurring within sequences, estimated values were excluded up until the completed presentation of the second PE, therefore corresponding to the first two (spatial task) and three (temporal task) sounds, respectively.

#### 6.10.2. Behavioral analysis

Individual accuracies, calculated as the percentage of correct trials per SAC level, were entered into a mixed-design repeated-measures ANOVA in MATLAB (The MathWorks Inc., 2023; version: 23.2.0 (R2023b), Natick, Massachusetts), with domain as a between-subjects factor and SAC level as a within-subjects factor. As Mauchly’s test indicated that the assumption of sphericity was violated (*W* = 0.024, *χ*^2^(9) = 158.01, *p* < .001), Greenhouse-Geisser corrected (ϵ = 0.36) degrees of freedom were used. The normality assumption was violated in both extreme conditions (Shapiro-Wilk-test: SAC = 1, *p* = 0.019; SAC = 5, *p* = 0.001), most strongly in SAC = 5, where accuracies approached ceiling. To fare on the safe side, a second mixed-design repeated-measures ANOVA was conducted only over the normally distributed SAC levels 2-4 to confirm significance.

We entered average confidence ratings per SAC level into an analogous mixed-design repeated-measures ANOVA. As Mauchly’s test indicated that the assumption of sphericity was violated (*W* = 0.22, *χ*^2^(9) = 64.46, *p* < .001), Greenhouse-Geisser corrected (ϵ = 0.64) degrees of freedom were used. Mauchly’s test confirmed normality in all conditions.

#### 6.10.3. Cluster-based permutation test

To add an additional analysis independent of the deconvolution-model a cluster-based permutation analysis in MATLAB (The MathWorks Inc., 2023; version: 23.2.0 (R2023b), Natick, Massachusetts) was performed, using the Fieldtrip toolbox (Version: 20250318) [94]. A “low surprisal” and a “high surprisal” condition was constructed by a median split of all surprisal values. All sections of pupil trace within a sequence (excluding first and last motion) where a defining “low surprisal” or “high surprisal” stimulus was preceded by two “low surprisal” sounds were binned according to the defining third stimulus. The resulting conditions featured a three-sounds sequence of either “low-low-low” or “low-low-high”, therefore starting with a mutual baseline before diverging with the defining third stimulus. Instantaneous pupil dilation rates, quantified as the difference between adjacent samples, were averaged per participant and condition. Due to the variable SOA in the temporal group, this analysis was only performed in the spatial group, where sounds from different trials and participants could be synchronized in large numbers. A cluster-based permutation analysis (dependent samples *t*-test) was performed over the duration of 0-2500 ms after onset of the defining third stimulus. Clustering was based on the maximum sum of *t*-values within a cluster. Significance was defined as a cluster-forming alpha of *p* = 0.05 at 1000 permutations.

#### 6.10.4. Cross-domain regression analysis for surprisal

A mixed effects linear regression analysis was conducted on the entire data over both experiments, using the *lmer()* function from the *lme4* package (Version: 1.1-37 [95]) in R (Version: 4.5.0)[96]. The models were subsequently compared to each other with a likelihood-ratio test, using a significance criterion of α = .05. The *full model* included in total nPar = 6 parameters: the fixed effects of (1) surprisal, (2) domain and (3) their interaction on pupil dilation with (4) an intercept, as well as (5) the variance for random slopes for surprisal across participants and (6) the residual variance. Random intercepts were removed [97] for the model to converge normally. This resulted in the following model equation for the full model:

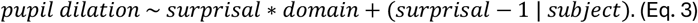

The *reduced model* only included the effects of surprisal and random slopes for surprisal across participants, eliminating the main effect for domain and its interaction with Surprisal (nPar = 4). The random intercepts were again removed, thus resulting in the formula:

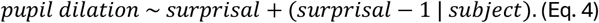

The two tasks were not considered as a random effect or grouping variable (but as a fixed effect) as they do not induce nonindependence [97], with participants being chosen randomly from the same population in both experiments. The models were fitted using maximum likelihood estimation, and Satterthwaite’s method was used to compute the degrees of freedom for the *t*-tests. Effect sizes are provided by *lme4* as R^2^ conditional (R^2^c) and marginal (R^2^m).

#### 6.10.5. Cross-domain regressions for infogain

To test whether *infogain* outperforms surprisal when predicting pupil dilation we performed a further model comparison, akin to the previously described procedure. Note that this marks the beginning of exploratory analyses that have not been preregistered, in contrast to the analyses presented up to now. As previously, infogain was Box-Cox transformed and z-scored before entering the regression analysis. Again, a *full model* included the effects of infogain, domain and their interaction on pupil dilation, grouped by participants, resulting in the formula:

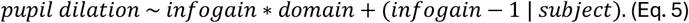

The *reduced model* only included the effects of the latent variable, grouped by participants, thus resulting in the formula:

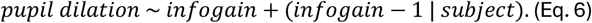

The full and the reduced model were compared via Bayes factor (BIC approximation from *lme4*) to choose the winning model for the latent variable infogain. The winning infogain model was compared against the winning surprisal model.

