## Supplement: Bayesian Model Description for "Arousal-related mediation of perceptual belief updating across auditory domains"

### Supplement: detailed Bayesian model description

#### 1. Generation of sensory signals

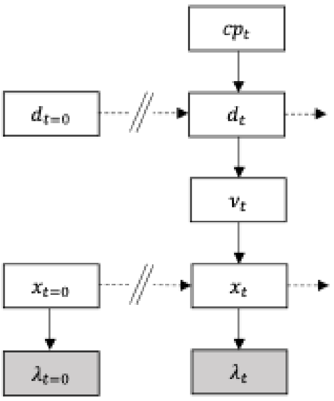


$$cp_{t=1} = 1$$

$$cp_{t>1} \sim 1B(1, H)$$

$$d_{t=0} \sim 2B(1{,}0.5) - 1$$

$$d_t = \begin{cases} d_{t-1} & if\ cp_t = 0 \\ -d_{t-1} & if\ cp_t = 1 \end{cases}$$

$$v_t \sim N\left(v_t;\ d_t\ \mu_{exp}, \sigma^2_{exp}\right)$$

$$x_{t=0} \sim U(-40°, 40°)$$

$$x_t \sim x_{t-1} + v_t$$

$$\lambda_t \sim N\left(\lambda_t;\ x_t,\ \left(\sigma_{sens,0} +\ \lambda_t|x_t|\ \right)^2\right)$$

We sampled the first sound location $x_{t=0}$ from a bounded uniform distribution between -40° and +40°. Subsequent sound locations $x_t$ were determined by velocity $v_t$ according to: $x_t = x_{t-1} + v_t$. At each timepoint $t \geq 1$, velocity $v_t$ was sampled at random from a normal distribution with variance $\sigma^2{}_{exp}$ and mean equal to $d_t * \mu_{exp}$. Here, $d_t \in \{-1, +1\}$ indicates the direction of the current movement direction, whereas $\sigma_{exp}$ and $\mu_{exp}$ determine the task difficulty. [We set them individually as $\mu_{exp} = 3 * MAA_{0°}$ and $\sigma_{exp} = 1 * M_{AA0°}$, where $MAA_{0°}$ is the minimum audible angle at 0° azimuth that was measured for every subject before starting the main task.] Movement direction $d_t$ was sampled at random at the beginning of each trial, but it would change suddenly during the sound sequence whenever a changepoint occurred: i.e. when $cp_t = 1$. Changepoints were sampled at random with a fixed hazard rate: $H = 1/5$. For analyses purposes, we defined every trial's first velocity, $v_t$=1, as having followed from a change point; i.e. $cp_{t=1} = 1$.

Observers only have access to noise-corrupted internal estimates $\lambda_t$ of the true sound locations $x_t$. We assume normally distributed sensory noise with standard deviation $\sigma_{sens}(x_t)$, which may linearly increase with the absolute stimulus location: $\sigma_{sens}(x_t) = \sigma_{sens,\,0°} + \gamma_\sigma|x_t|$, with $\gamma_\sigma \geq 0$.

#### 2. Prior knowledge and likelihood functions

Participants were superficially informed by the experimenter about the concepts of changepoints and experimental noise, without mentioning specific values for hazard rate $H$ or experimental parameters $\mu_{exp}$ and $\sigma_{exp}$. Nevertheless, we assume that our observers learned about the velocity distribution, $N(v_t;\ \pm\mu_{exp},\ \sigma^2{}_{exp})$, during the main task practice block. We also assume that they were able to form a stable, but not necessarily accurate estimate of the constant hazard rate $H$. Moreover, we assume that observers have access to the instantaneous uncertainty of their own internal estimates, $\sigma_{sens}(\lambda_t) = \sigma_{sens}(x_t) = \sigma_{sens},0° + \gamma_\sigma|x_t|$. Hence, their likelihood function for sound location $x_t$ is normally distributed and centred on internal estimate $\lambda_t$:

$$p(\lambda_t|x_t,) = N(x_t, \lambda_t, \sigma^2_{sens}\ (\lambda_t)) \qquad \text{(Eq. 1)}$$

Combining equation 1 and $v_t = x_t - x_{t-1}$, it follows that the likelihood function for velocity $v_t$ must also be normally distributed, with mean equal to the difference of the last two consecutive internal estimates, and variance equal to the sum of the variance for each stimulus location:

$$p(\lambda_t - \lambda_{t-1}|v_t,) = N(v_t, \lambda_t - \lambda_{t-1}, \sigma^2_{sens}\ (\lambda_t) + \sigma^2_{sens}\ (\lambda_{t-1})) \qquad \text{(Eq. 2)}$$

Furthermore, since velocity $v_t$ was sampled from a normal distribution according to the generative process, $v_t \sim N(v_t;\ d_t\mu_{exp},\ \sigma^2{}_{exp})$, we can compute the likelihood of observing $\lambda_t - \lambda_{t-1}$ conditional on either direction as:

$$p(\lambda_t - \lambda_{t-1}|d_t,) = N\left(\lambda_t - \lambda_{t-1};\ d_t\mu_{exp}\ ,\ \sigma^2_{exp} + \sigma^2_{sens}\ (\lambda_t) + \sigma^2_{sens}\ (\lambda_{t-1})\right) \qquad \text{(Eq. 3)}$$

In other words, the likelihood for a left-/rightward direction $d_t$ can be computed by evaluating the probability density of $N\left(\pm\mu_{exp}, \sigma^2_{exp} + \sigma^2_{sens}\ (\lambda_t) + \sigma^2_{sens}\ (\lambda_{t-1})\right)$ at the point $\lambda_t - \lambda_{t-1}$.

### 3. Bayesian directions Change Point (BdCP) model

An observer's task is to infer the direction of the sound movement at the end of the sequence. Thesimplest estimate of $d_t$ is given by the likelihood ratio:

$$LikeRatio_{d_t} = \frac{p(\lambda_t - \lambda_{t-1}|d_t = 1)}{p(\lambda_t - \lambda_{t-1}|d_t = -1)} = \frac{LikeL_t}{LikeR_t} \quad \text{(Eq. 4)}$$

If this ratio is larger than one, then a leftward movement is more likely. Vice versa, a $LikeRatio_{d_t}$ < 1 indicates that a rightward movement is more likely. This is called the maximum likelihood method. However, such a strategy that bases its direction responses merely on the relative likelihood for either direction would give many erroneous responses if there were much sensory noise. I.e. when $\sigma^2{}_{sens}(\lambda_t)$ + $\sigma^2{}_{sens}(\lambda_{t-1})$ is large it is likely that the instantaneous internal estimate of the current velocity, $\lambda_t$ − $\lambda_{t-1}$, is wildly inaccurate and a change of sign may frequently occur. Instead, Bayesian theory postulates that ideal observers can minimize their perceptual errors by making use of all available sensory information ($\lambda_{1:t}$) and their knowledge of the generative process for these sensory signals, including learned parameters $\mu_{exp}$, $\sigma_{exp}$, and $H$. In particular, one can make use of the fact that changepoints only occur sometimes. This means that a belief about the previous movement direction is also useful information about the current movement direction. According to the Bayesian belief updating framework, such pre-existing beliefs are used to construct a "prior". After hearing a new sound, the prior information is then integrated with the information from the latest likelihood function to obtain a "posterior".

At the start of a sequence, each direction is equally likely (50%). The prior ratio thus equals 1:

$$PriorRatio_{d_{t=1}} = \frac{p(d_{t=1}=1)}{p(d_{t=1}=-1)} = \frac{PriorL_t}{PriorR_t} = 1 \quad \text{(Eq. 5)}$$

After hearing a new sound, the likelihood ratio is computed according to equation 4. Subsequently, prior and likelihood are integrated by multiplication to form a posterior ratio:

$$PostRatio_{d_t} = \frac{p(d_{t=1} = 1|\lambda_{0:t})}{p(d_{t=1} = -1|\lambda_{0:t})} = \frac{PostL_t}{PostR_t} = \frac{c^{-1} * PriorL_t * LikeL_t}{c^{-1} * PriorR_t * LikeR_t} = PriorRatio_{d_t} * LikeRatio_{d_t} \quad \text{(Eq. 6)}$$

Notice that the normalization constant from Bayes' rule, $c = PriorL_t * LikeL_t + PriorR_t * LikeR_t$, drops out and does not need to be computed. To form the next prior ratio, the posterior ratio is updated to account for the possibility of a changepoint before the presentation of the next sound:

$$PriorRatio_{d_{t>1}} = \frac{p\left(d_{t=1} = 1\middle|\lambda_{0:(t-1)}\right)}{p\left(d_{t=1} = -1\middle|\lambda_{0:(t-1)}\right)} = \frac{PriorL_t}{PriorR_t} = \frac{(1-H)PostL_{t-1} + HPostR_{t-1}}{(1-H)PostR_{t-1} + HPostL_{t-1}} \quad \text{(Eq. 7)}$$

So, a Bayesian ideal observer model iteratively updates its beliefs by following equations 4, 6 and 7. After the last sound has played, the final decision is made based on the posterior ratio (equation 6).
